# Polymer brush inspired by ribosomal RNA transcription

**DOI:** 10.1101/2023.03.16.533038

**Authors:** Tetsuya Yamamoto, Wei Li

## Abstract

Pre-ribosomal RNAs are synthesized during the transcription by RNA polymerase I molecules localized at the surfaces of a nucleolus subcompartment. Inspired by the ribosomal RNA transcription, we here develop a scaling theory of a brush of polymers, where monomers are added to their grafted ends in the steady state. Our theory predicts that monomers newly added to the polymers stay at the vicinity of the surface due to the slow dynamics of the polymers and thus the polymer volume fraction increases with increasing the polymerization rate. The excluded volume interaction between polymers and reactant monomers suppresses the diffusion of reactant monomers and thus decreases the polymerization rate. The extent of the suppression of monomer diffusion increases with increasing the polymerization rate because the diffusion length decreases, rather than the condensation of polymers due to their slow dynamics.

## 1 Introduction

Nucleolus is a condensate of ribosomal RNAs and proteins and is a factory of ribosomes. In higher organisms, nucleolus forms a multi-phase structure, where multiple subcompartments, called fibrillar centers (FCs), are dispersed in another subcompartment, called a granular component (GC) [1], see fig. 1**a**. Indeed, distinct phases, called dense fibrillar components (DFCs), are assembled between each FC and GC. Nucleoli and their subcompartments have been thought to be assembled by liquid-liquid phase separation (LLPS) because of the fact that the major protein components of each phase show phase separation in test tube experiments [2]. However, condensates assembled by LLPS show coarsening and coalescence to minimize the surface energy. It is in contrast to the situation of nucleolus, in which multiple FCs are dispersed.

**Fig. 1.**
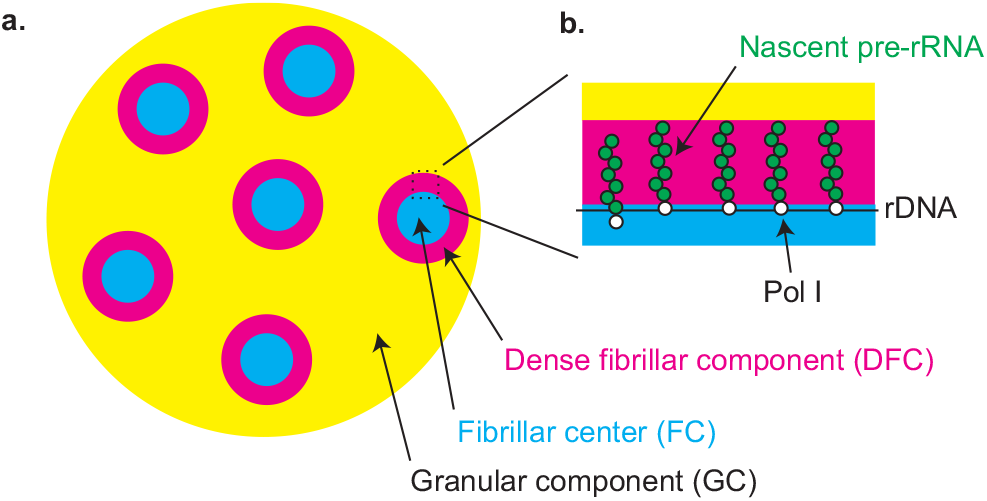
Multi-phase structure of nucleolus: **a.** A nucleolus is composed of three phases – fibrillar centers (FCs), granullar component (GC), and dense fibrillar component (DFC). Multiple FCs are dispersed in a GC and there is a layer of DFC at the surface of each FC. **b.** Ribosomal DNAs (rDNAs) are localized at the surfaces of FCs and is transcribed by RNA polymerase Is (Pol Is) entrapped in FCs. Nascent pre-rRNAs are end-grafted to the surfaces of FCs via Pol Is. These nascent pre-rRNAs are modeled as a polymer brush.

Indeed, if RNA polymerase I (Pol I), which transcribes precursor ribosomal RNAs (pre-rRNAs), is depleted, FCs show fusion to be one large condensate, much like condensates assembled by LLPS [3]. This implies that the transcription of pre-rRNAs somehow suppresses the fusion of FCs. The transcription of pre-rRNAs happens at the interface between FCs and DFCs [4, 5]. Multiple copies of pre-rRNAs are tandemly repeated in the DNA region localized at the interfaces (only 2 - 3 active copies per FC in an estimate [4], but can be more) and many Pol Is are loaded to each active copy of length ~ 15 kbps for transcription [1]. One interesting hypothesis is that nascent pre-rRNAs, which are end-grafted to the surfaces of FCs via Pol Is, act as polymer brushes that suppress the fusion of FCs [6], much like the situation that polymer brushes at the surfaces of colloids suppress the flocculation as predicted by Witten and Pincus [7], see fig. 1**b**.

Single-stranded RNAs are relatively flexible polymers with the aspect ratio of its Kuhn unit in the order of unity. In the physiological salt concentration, the electrostatic interaction between RNAs is screened by salt ions. Nascent pre-rRNAs end-grafted at a surface can be thus treated as a brush of electrically neutral flexible polymers. The simplest model of a neutral flexible polymer brush is the Alexander model, with which the monomer concentration is assumed to be uniform in the brush [8]. By using the Flory-type argument, Alexander predicted that the height of the brush is proportional to the number of monomers in the chains, implying that the chains in the brush are stretched due to the excluded volume interaction with the neighboring chains. The scaling argument by de Gennes augments the structures in the length scales shorter than the brush height by using the fact that the correlation length of chains in the brush is set by the distance between the grafting points [9]. The self-consistent field theory predicts that the monomer concentration is a parabolic function of the distance from the surface in the classical limit [10–12].

In general salt conditions, RNAs should be treated as polyelectrolytes. In the low salt concentration, the charges of polyelectrolytes in the brush are mainly neutralized by counterions. In his pioneering work, Pincus used the scaling theory to predict that if the linear charge density and the grafting density of polyelectrolytes in the brush are large enough, counterions are entrapped in the brush region due to the strong electrostatic interaction between the polyelectrolytes and counterions, and the polyelectrolytes are swollen due to the osmotic pressure of the counterions (osmotic regime) [13]. If the free energy due to the translational entropy of counterions dominates the free energy due to the electrostatic interaction between polyelectrolytes and counterions, the brush collapses because counterions escape from the brush (Pincus brush regime). Polyelectrolyte brushes experience the crossover between the osmotic and Pincus brush regimes by applying electric fields because polyelectrolytes collapse to neutralize the charges induced by the applied electric fields [14]. In the high salt concentration, polyelectrolyte charges are mainly neutralized by salt ions (salty brush regime). In this regime, a polyelectrolyte brush behaves as a brush of electrically neutral polymers because the monomer-monomer electrostatic interactions are screened by salt ions and are reduced to the second virial coefficient [13].

Chemically similar to a polymer brush of RNAs is a polymer brush of DNA. DNA brushes are useful systems to study how the transcription dynamics depends on the local packing density of DNA [15–17]. In the low salt concentration, DNAs in a brush are swollen by the osmotic pressure of counterions due to the polyelectrolyte nature of DNA [18]. Double-stranded DNA is a semiflexible polymer with the aspect ratio of its Kuhn unit is in the order of 100 [19]. Schaefer, Joanny, and Pincus predicted that semiflexible polymers have the marginal solvent regime, in which the mean-field theory is effective although the two-body interaction between the Kuhn segments dominates the three-body interaction between them, because the excluded volume of segments with high aspect ratio is small [20]. In the physiological salt concentration, DNA brush behaves as a brush of electrically neutral semiflexible polymers in the marginal solvent regime [18]. RNA brushes probably do not have the marginal solvent regime because the aspect ratio of their Kuhn segments is relatively small.

What is the difference between the nascent pre-rRNA brushes at the surfaces of nucleolus subcompartments and the above-mentioned concepts of polymer brushes? First, RNA-binding proteins (RBPs) are bound to some RNAs and the multi-valent interaction between these RBPs drives LLPS [21, 22]. Indeed, nuclear bodies, such as paraspeckles, are assembled by the multi-valent interaction between RBPs bound to architectural RNAs [22–25]. Fibrillarin RNA-binding proteins are a major component of the DFC layers at the interface between FCs and GC and are the RBPs of nascent pre-rRNAs [2, 4]. Fibrillarin shows phase separation in the physiological concentration [2]. In our recent theory, we have predicted that the DFCs at the interface between FCs and GC are polymer brushes of nascent pre-rRNAs that are swollen to accommodate RBPs in the brush [6]. There are analogous systems with DNA – DNA brushes with DNA-binding proteins, such as histones [26, 27], transcription factors [28], and Structural Maintenance of Chromosome proteins [29–32]. Second, nascent pre-rRNAs are end-grafted to the surface via Pol Is that add nucleoside triphosphates (which are monomers of RNAs) to the nascent pre-rRNAs. Because of the slow dynamics of polymers, RNA monomers newly added to the nascent pre-rRNAs stay at the vicinity of the surface for a relatively long time. The high concentration of RNA monomers at the surface may suppress the diffusion of reactant RNA monomers and limit the kinetics of transcription. This situation is very different from cases in which monomers are added to the free ends of polymers in a brush, such as the free-radical polymerization initiated from a surface [33, 34].

Inspired by the dynamical nature of the nascent pre-rRNA brush, we here use the scaling theory to predict the structure of polymers in a brush, where monomers are added to the grafted-ends with a constant rate (sec. 2) and the polymerization rate, which is due to the diffusion of reactant monomers (sec. 3). We do not take into account the transcriptional termination and co-transcriptional processing to focus on the polymer dynamics driven by the polymerization reaction. The nascent pre-rRNAs during transcription is a type of living polymers, which are defined as non-equilibrium polymers due to the polymerization and depolymerization kinetics [35]. Indeed, our theory predicts that the elongation of Pol I transcription is too slow for the polymer dynamics to be relevant for the case of pre-rRNA transcription. However, we anticipate that this theory can be extended to polymerization reaction in other biological contexts and may motivate polymer chemists to design polymer brushes by taking advantage of the slow relaxation dynamics of polymers.

## 2 Polymer brush inspired by pre-rRNA transcription

### 2.1 Mushroom regime

We first treat a single polymer (such as nascent pre-rRNA) end-grafted to a planar surface via a catalyst (such as Pol I) that adds monomers to its grafted end with a constant rate *k*_p_ in the steady state. For simplicity, we treat the dynamics of polymers in the brush by using the Rouse model, with which the hydrodynamic interaction between monomers is neglected [36, 37]. With this model, the diffusion constant of a subchain composed of *g* monomers is *D_g_* (= *D*_1_/*g*), where *D*_1_ is the diffusion constant of a monomer. The size of the subchain is *ξ* = *bg*^3/5^ in an athermal solvent with the Flory exponent. The relaxation time of the subchain is *ξ*^2^/*D_g_* (= *τ*_1*g*_^11/5^), where *τ*_1_ (= *b*^2^/*D*_1_) is the relaxation time of a monomer. The number *g*(*t*) of monomers in a subchain that are relaxed in time *t* is

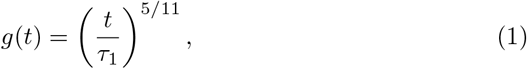

where it is derived by *ξ*^2^/*D_g_* ≈ *t*.

The number of monomers newly added to the chain in time *t* is *k*_p_*t*. The relaxation is faster than the polymerization for *k*_p_*t* < *g*(*t*). The polymerization kinetics becomes as fast as the relaxation at a crossover time 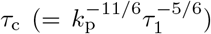. The number of monomers in a subchain relaxed in the time *τ*_c_ is

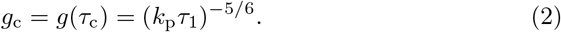

The size of the subchain is

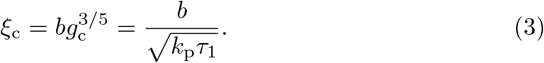

The polymer volume fraction at this length scale is

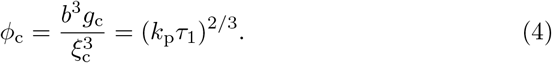

Now we consider the structure of the chain in the length scale larger than *ξ_c_*, see fig. 2**a**. *ξ_c_* is the minimum length scale of the relaxed region in the chain. Eq. (1) predicts that the subchain composed of *g*(*t* – *t*_0_) monomers around the monomer polymerized at time *t*_0_ is already relaxed at time *t* (*t* – *t*_0_ > *τ_c_*). The polymers are therefore considered as the chains of relaxed blobs of varying sizes, see fig. 2**a**. The chain is not relaxed at the length scale larger than a relaxed blob *ξ*(*t*–*t*_0_). The relaxed blobs thus stay at the position of the catalyst, while they are pushed out by the blobs of subchains of newly synthesized monomers, see fig. 2**a**. The relaxed blobs are thus packed around the catalyst. The volume *r*^3^ occupied by the subchain that is polymerized in time *t* (> *τ_c_*) has the form

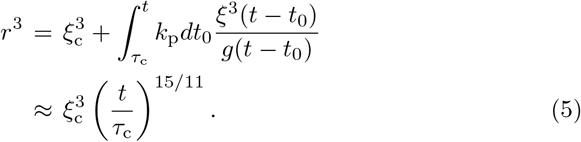

**Fig. 2.**
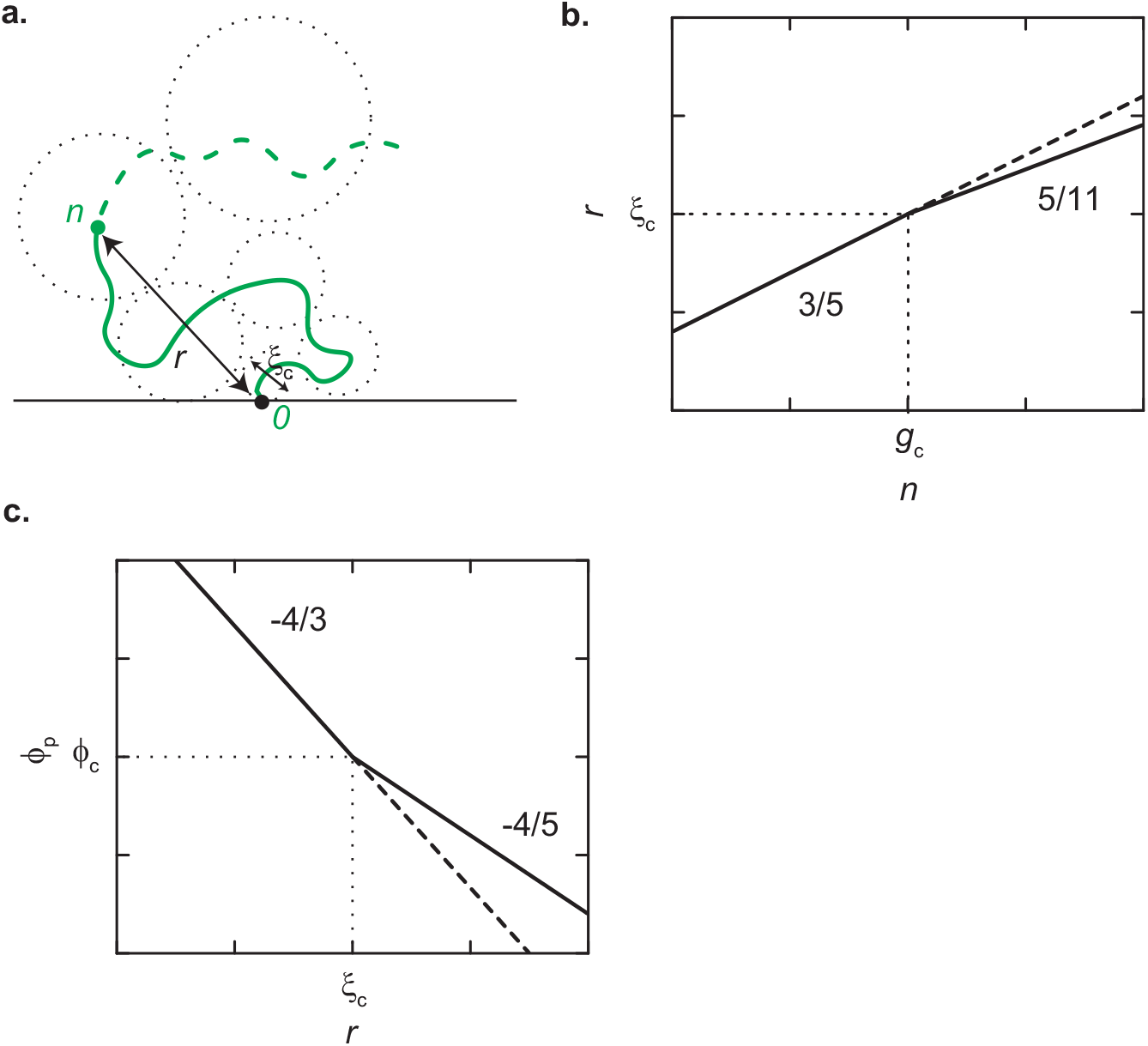
Mushroom regime. **a.** A polymer (nascent pre-rRNA) synthesized by a catalyst (Pol I) at a surface. In general, there can be multiple catalysts at the surface, but they are separated so that the interaction between end-grafted polymers is negligible (mushroom regime). Monomers are added to the grafted-end of the polymer with a constant rate. The polymer is considered as a chain of blobs, where the subchain inside of each blob is already relaxed. The smallest blob has the length *ξ*_c_ and is composed of *g*_c_ monomers, see eqs. (2) and (3). These blobs are space-filling. **b.** The size *r* of a subchain is shown as a function of the number *n* of monomers of the subchain (double-logarithm plot). **c.** The polymer volume fraction *ϕ*_p_ is shown as a function of the distance *r* from the grafting point (doublelogarithm plot). The broken lines in **b** and **c** are the case of an end-grafted polymer in the thermodynamic equilibrium.

The subchain is composed of *n* (= *k*_p_*t*) monomers and thus the size of the subchain is

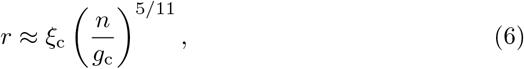

see fig. 2**b**. This approach is an extension of the Daoud-Cotton scaling theory [38] to the relaxed blobs filled in the geometry of a planer brush. The local polymer volume fraction for *r* > *ξ*_c_ thus has the form

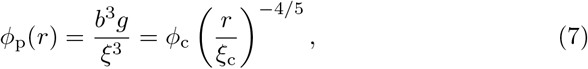

see fig. 2**c**. Eq. (7) is derived by using *ξ* = *bξ*^3/5^ and then use eq. (1), *n* = *k*_p_*t*, and eq. (6) to represent it as a function of *r*. Eq. (7) indeed has the same form as the average polymer volume fraction *b*^3^*n/r*. The polymer volume fraction *ϕ*_p_ thus is larger than an end-grafted polymer at the equilibrium for *r* > *ξ*_c_ due to the slow relaxation dynamics of polymers, compare the solid and broken lines in fig. 2**c**.

### 2.2 Brush regime

Next, we treat a brush of polymers that are end-grafted to a surface via catalysts that add monomers to their grafted-ends with a constant rate *k*^p^, see fig. 3**a**. These catalysts are distributed to the surface with the surface density *σ*. The correlation length of the brush is the distance *ξ*_b_ (= *σ*^-1/2^) between the grafting points [9]. In the length scale smaller than *ξ*_b_, the polymer conformation follows the same scaling relationship with the case of mushroom regime because there is no interaction between polymers, see the region of *z* < *ξ*_b_ in fig. 3**b**. By using eq. (6), the number of monomers in *r* < *ξ*_b_ is derived as

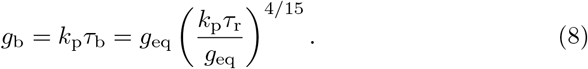

**Fig. 3.**
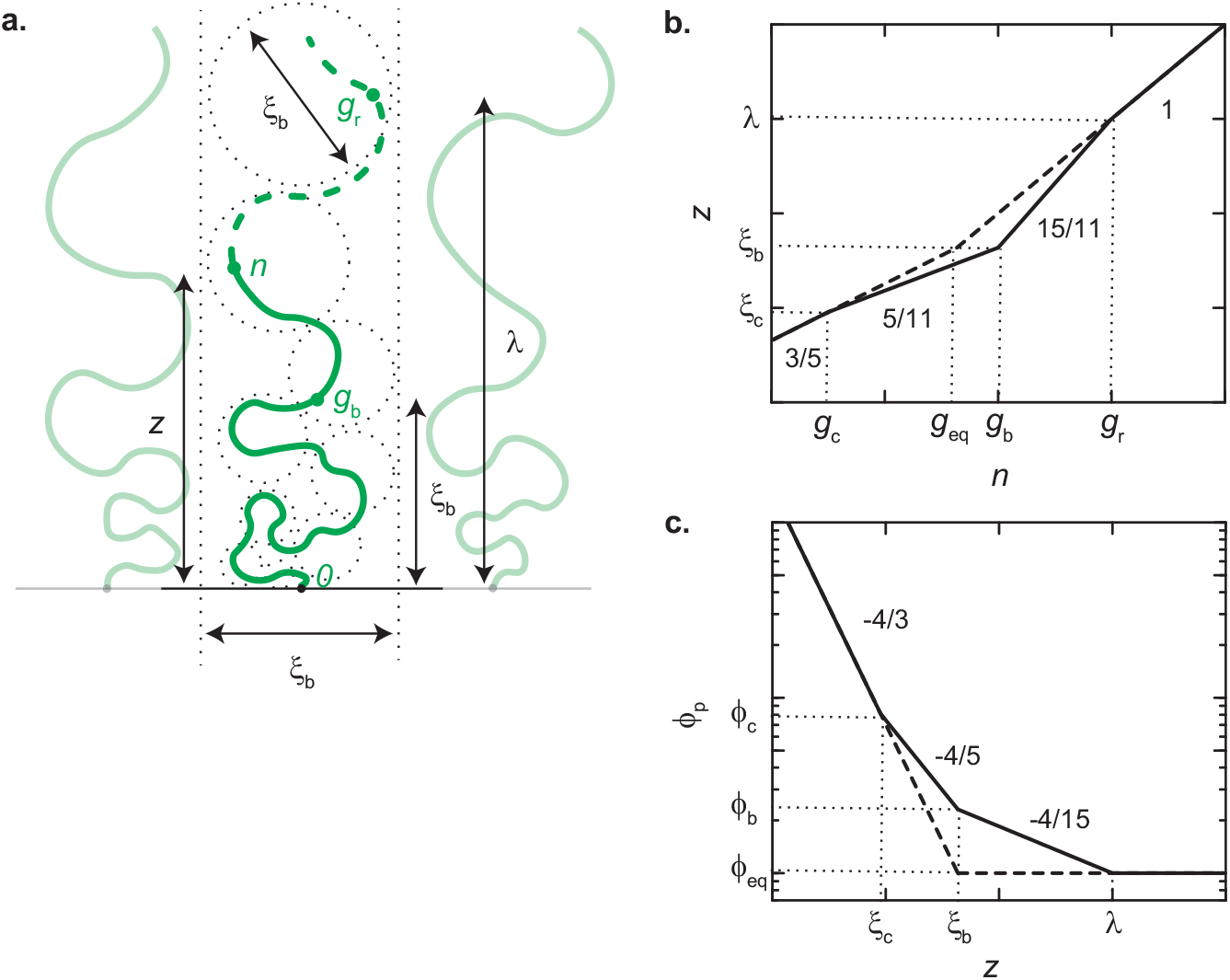
Brush regime: **a.** Polymers (pre-rRNAs) synthesized by catalysts (Pol Is) at a surface. The grafting density *σ* is relatively large so that the interaction between the end-grafted polymers is significant. Monomers are added to the grafted-ends of the polymers with a constant rate (brush regime). An end-grafted polymer in the brush is considered as a chain of blobs, where the subchain in each blob is already relaxed. These blobs are space-filling and are confined in the area 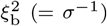 due to the interaction between end-grafted polymers. **b.** The height *z* of a subchain is shown as a function of the number *n* of monomers of the subchain (double-logarithm). **c.** The polymer volume fraction *ϕ*_p_ is shown as a function of the distance *r* from the surface (double-logarithm). The broken lines in **b** and **c** are the case of an end-grafted polymer in the thermodynamic equilibrium.

In eq. (8), we used the number *g*_eq_ of monomers in the correlation volume 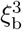 in the thermodynamic equilibrium and the relaxation time *τ*_r_ of the subchain composed of *g*_eq_ monomers:

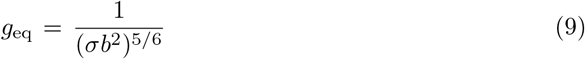

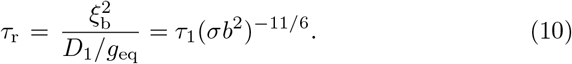

The number *g*_b_ of monomers in the volume 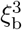 returns to that *g*_eq_ in the equilibrium for 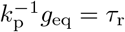, see eq. (8). The number of monomers in the region *r* < *ξ*_b_ increases with increasing the polymerization rate *k*_p_. The polymer volume fraction at the length scale of *ξ*_b_ is 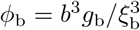, see eq. (8).

In the length scale larger than *ξ*_b_, the relaxed blobs are confined in the area 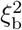 due to the interaction between end-grafted polymers and are stacked towards the *z*-direction. The height *z* of *n*-th monomer from the grafted-end is

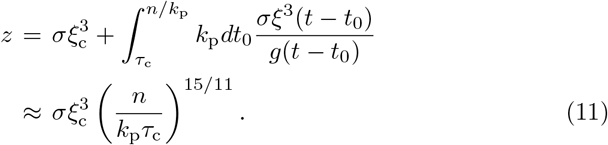

Note that *z* = *ξ*_b_ for *n* = *g*_b_ and thus eq. (11) smoothly crossovers with eq. (6) at *n* = *g*_b_.

The size of the relaxed blob *ξ*(*t*) becomes the size of the blob *ξ*_b_ of the equilibrium brush in the time *t* = *τ*_r_. The distance with which a monomer travels in the relaxation time *τ*_r_ is derived by substituting *n* = *g*_r_ (= *k*_p_*τ*_r_) into eq. (11) as

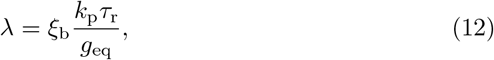

see fig. 3**b**. It is interesting to note that eq. (12) has the form of the brush height of polymers composed of *g*_r_ (= *k*_p_*τ*_r_) monomers in the thermodynamic equilibrium. This implies that the effect of the slow dynamics of polymers is not dramatic. Eq. (12) returns to *λ* = *ξ*_b_ for 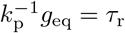.

The local polymer volume fraction for *ξ*_b_ < *z* < *λ* has the form

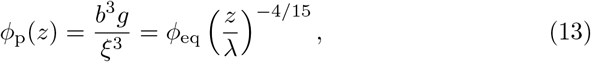

see fig. 3**c**. Eq. (13) is derived by using *ξ* = *bg*^3/5^ and then use eq. (1), *n* = *k*_p_*t*, and eq. (11) to represent it as a function of *z*. Eq. (13) has the same form as the average polymer volume fraction *b*^3^*σn/z*. In eq. (13), we used the polymer volume fraction *ϕ*_eq_ of a blob of the equilibrium brush

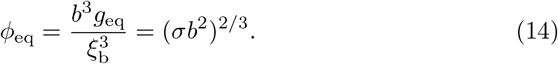

The polymer volume fraction *ϕ*_p_(*z*) for *z* > *λ* is *ϕ*_eq_, see fig. 3**c**.

### 2.3 Brush height

For the case of the transcription of nascent pre-rRNAs on the surfaces of FCs, these pre-rRNAs experience the co-transcriptional processing and are released when Pol Is reach the terminator sequence. To focus on the roles played by the polymerization kinetics in the structure of the polymer brush, we neglect these complex processes and consider cases in which the end-grafted polymers are randomly scissored by other catalysts (such as ribonucleases). In this case, the number *N* of monomers in each end-grafted polymer in the brush follows the kinetic equation

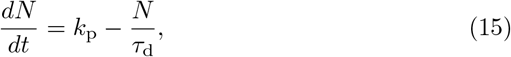

where *τ*_d_ is the average time of polymer degradation. In the steady state, the number of monomers in each end-grafted chain is *N* = *k*_p_*τ*_d_.

i. The case of *τ*_d_ > *τ*_r_: In the limit of small polymerization rate *k*_p_, the number *N* of monomers in each end-grafted polymer is too small and the interaction between the polymers is negligible (mushroom regime). Each end-grafted polymer is relaxed in the entire length scale for *k*_p_*τ*_r_/*g*_eq_ < 1 (qusi-equilibrium mushroom regime) and thus the size of the end-grafted polymer is

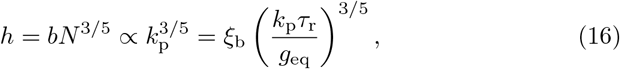

see the cyan line in fig. 4. It is not the true equilibrium because of the polymerization and degradation. For *h* > *ξ*_b_, the interaction between end-grafted polymers becomes significant and the end-grafted polymers thus become a brush (quasi-equilibrium brush regime). The brush height has the form

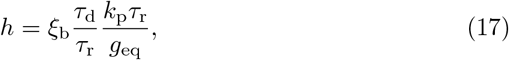

see the cyan line in fig. 4. For the case of *k*_p_*τ*_r_/*g*_eq_ < 1, the polymers in the brush are relaxed in the entire length scale. In contrast, the polymer volume fraction becomes larger than that at the equilibrium in the region *ξ*_c_ < *z* < *λ*, see fig. 3. The brush height is given by eq. (17) both for *k*_p_*τ*_r_/*g*_eq_ < 1 and *k*_p_*τ*_r_/*g*_eq_ > 1. This implies that although the polymer volume fraction increases due to the slow polymer dynamics in *ξ*_c_ < *z* < *λ*, its effect on the brush height is only within the numerical factor of order unity, see also fig. 3**b**.
ii. The case of *τ*_d_ < *τ*_r_: In the limit of small polymerization rate k_p_, the system is in the quasi-equilibrium mushroom regime, see eq. (16) and the magenta line in fig. 4. For *k*_p_*τ*_r_/*g*_eq_ > (*τ*_r_/*τ*_d_)^6/11^, the end-grafted polymers are relaxed in the length scales smaller than *ξ*_c_ (more precisely, the size of a relaxed blob), but not in the larger length scale (non-equilibrium mushroom regime). The size of the non-equilibrium mushroom is

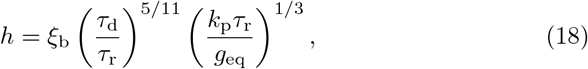

see the magenta line in fig. 4. Eq. (18) is derived by substituting *n* = *N* into eq. (6). For *h* > *ξ*_b_, the interaction between end-grafted polymers becomes significant (non-equilibrium brush regime). The brush height is

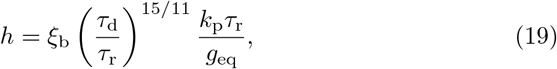

see the magenta line in fig. 4. Eq. (19) is derived by substituting *n* = *N* in eq. (11). In the non-equilibrium brush regime, the end-grafted polymers are relaxed in the length scale smaller than *ξ*_c_ (more precisely, the size of a relaxed blob), but not in the larger length scale.

**Fig. 4.**
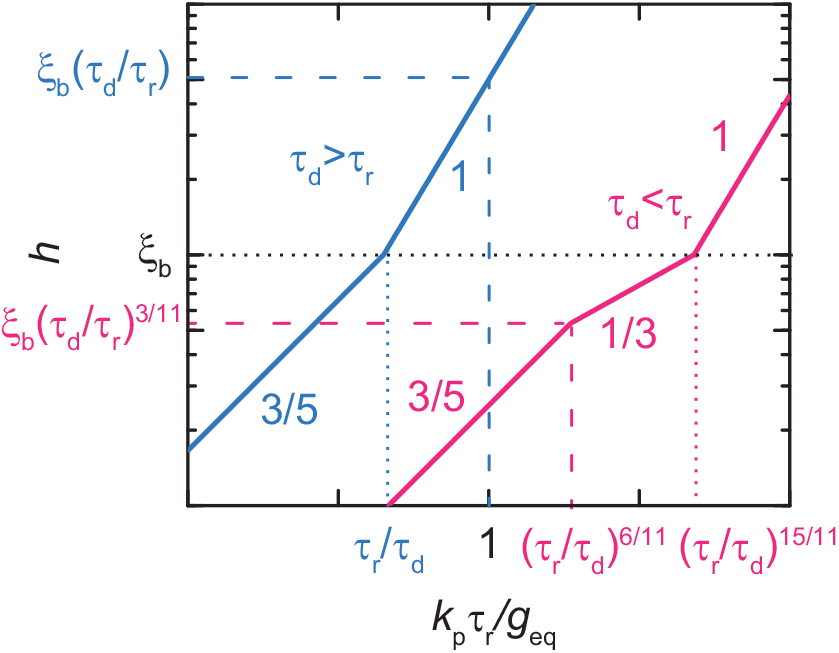
Brush height: The brush height *h* is shown as a function of rescaled polymerization rate *k*_p_*τ*_r_/*g*_eq_ for *τ*_d_ > *τ*_r_ (cyan) and *τ*_d_ < *τ*_r_ (magenta). *τ*_d_ is the degradation time of polymers and *τ*_r_ is the relaxation time, see eqs. (10) and (15). The end-grafted polymers form mushrooms for *h* < *ξ*_b_ and form a brush for *h* > *ξ*_b_, see the black dotted line. The crossovers between the mushroom and brush regimes are shown by the cyan and magenta dotted lines. The cyan and magenta broken lines are the crossover between the quasi-equilibrium and non-equilibrium regimes.

## 3 Polymerization kinetics

### 3.1 Dynamics of reactant monomers

In the previous section, we derived the polymer volume fraction and the brush height by treating the polymerization rate *k*_p_ as a control parameter. The polymerization rate *k*_p_ depends on the volume fraction of (unreacted) reactant monomers at the position of the catalysts and the flux of these monomers, where both of them are affected by the polymer volume fraction in the brush. In this section, we theoretically predict how the polymer dynamics driven by the polymerization affects the kinetics of polymerization. For simplicity, we treat cases in which the polymerization reaction is diffusion-limited.

The flux of reactant monomers has the form

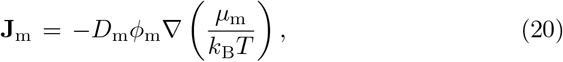

where *μ*_m_ is the chemical potential of reactant monomers

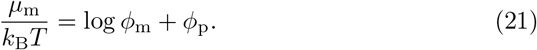

The flux **J**_m_ is defined as the volume of reactant monomers passing a unit area in a unit time and thus has the unit of m/s. Eq. (21) is derived by assuming that the solvent is athermal both for reactant monomers and monomers in polymers because of the same chemical structure, see also the Appendix A for the derivation.

We use the coordinate *z* to represent the positions in the brush for *z* > *ξ*_b_ and use the radial coordinate *r* to represent the position for *r* < *ξ*_b_. The catalyst acts as a sink of reactant monomers. In the steady state, the flux of reactant monomers in the *z*-direction is

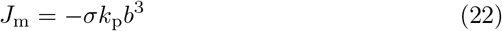

for *z* > *ξ*_b_. For *r* < *ξ*_b_, we use the flux in the radial direction

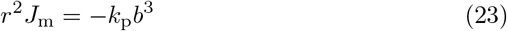

to ensure the continuity of the flux at *r* = *ξ*_b_. Eqs. (22) and (23) are formulated with the picture that the flux has the spherical symmetry in the length scale of the reaction radius and is approximated as the planar symmetry in the length scale larger than *ξ*_b_ because the length scale *ξ*_b_ is much larger than the reaction radius rc (much like the fact that electromagnetic waves are assumed to be planar at far away from a point source [39]). We thus used the same notation *J_m_* for the radical component of the flux in *r* < *ξ*_b_ and the *z*-component of the flux in *z* > *ξ*_b_. The monomer volume fraction *ϕ*_m_ is derived by using eqs. (22) and (23) with eqs. (21) and (20), see also the appendix B for the general solution.

### 3.2 Boundary conditions

The top of the brush is in equilibrium with the external solution that acts as the reservoir of reactant monomers. Rigorously speaking, there is no steady state in the solution of the diffusion equation for the planar geometry. We thus assume that the polymer brush is on a spherical surface, but the surface can be approximated as a planar surface if its curvature is relatively small. The monomer volume fraction at the top of the brush, *z* = *h*, is

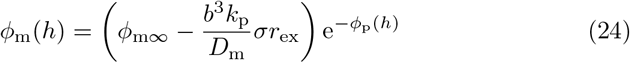

due to the local equilibrium at the brush top, where *r*_ex_ is the distance between the top of the brush and the center of the curvature of the spherical surface. The second term in the round bracket of eq. (24) is derived by assuming that the monomer volume fraction at the exterior of the brush follows the diffusion equation.

The length scale with which reactant monomers can diffuse in the reaction time 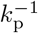 is 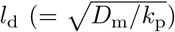. The diffusion constant of reactant monomers is approximately equal to the diffusion constant of monomers in the polymers, *D*_m_ ≈ *D*_1_, because both of the monomers have the same size. The diffusion length *l*_d_ is thus equal to the length scale *ξ*_c_, see eq. (3). This implies that the distribution of reactant monomers is at the local equilibrium for *r* < *ξ*_c_. If the polymerization reaction is diffusion-limited, the boundary condition reads

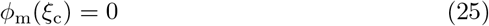

for cases in which the length scale *ξ*_c_ is larger than the reaction radius *r*_c_.

### 3.3 Perturbation analysis

The polymerization rate *k*_p_ is derived by using the relationship

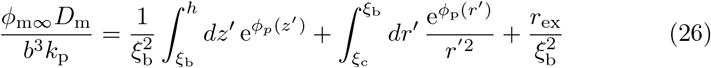

in the brush regime. Eq. (26) is derived by imposing the boundary conditions, eqs. (24) and (25), to the solutions of eqs. (22) and (23). The corresponding relationship for the mushroom regime is derived by replacing *ξ*_b_ to *h*. The subchain in a relaxed blob is swollen due to the two-body excluded volume interaction between the monomers. The polymer volume fraction *ϕ*_p_ is small in the length scale larger than *ξ*_c_ and eq. (26) can be expanded in a power series of *ϕ*_p_. We therefore treat the polymer volume fraction *ϕ*_p_ as perturbation and derive the polymerization rate *k*_p_ in the form

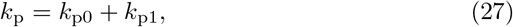

where *k*_p0_ is the polymerization rate for the case of *ϕ*_p_ = 0 and *k*_p1_ is an order linear to *ϕ*_p_.

For the case of *ϕ*_p_ → 0, eq. (26) is rewritten as

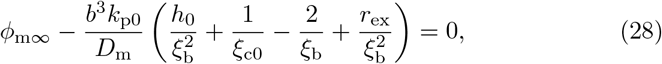

where *h*_0_ and *ξ*_c0_ are the brush height *h* and length scale *ξ*_c_ with the polymerization rate *k*_p0_. In the brush regime, *k*_p1_ is derived as

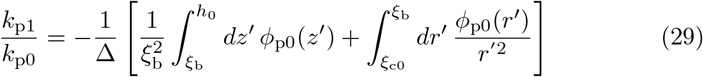

with

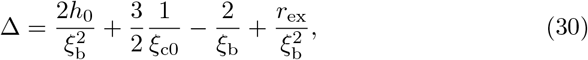

where *ϕ*_p0_ is the polymer volume fraction *ϕ*_p_ with the polymerization rate *k*_p0_. Eq. (29) is derived by expanding eq. (26) in a power series of *k*_p1_ and *ϕ*_p_ and omitting the higher order terms. The corresponding relationship for the mushroom regime is derived by replacing *ξ*_b_ to *h* (more precisely, Δ has an additional term –2/(3*h*_0_) in the mushroom regime, but it is smaller than other terms and is omitted). Because *r*_ex_ is the largest length scale in eq. (30), we use 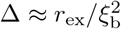 in the following.

1. The case of *τ*_d_ > *τ*_r_: The end-grafted polymers form a brush at the crossover between the quasi-equilibrium and non-equilibrium regimes, *k*_p0_*τ*_r_/*g*_eq_ = 1, see the cyan line in fig. 4. For *k*_p0_*τ*_r_/*g*_eq_ < (*τ*_r_/*τ*_d_)^2/3^, the distribution of reactant monomers in the entire region of the brush (or the mushrooms) is at the thermodynamic equilibrium, *l*_d_ > *h*_0_, and thus the polymer volume fraction in the brush (or mushroom) does not affect the polymerization rate, *k*_p1_ = 0, see the cyan solid line in fig. 5**a**. For (*τ*_r_/*τ*_d_)^2/3^ < *k*_p0_*τ*_r_/*g*_eq_ < 1, the distribution of reactant monomers in the brush (or the mushrooms) become out of the equilibrium, *l*_d_ < *h*_0_, but the polymers are relaxed in the entire length scales. The form of the deviation *k*_p1_ of the polymerization rate due to the polymer volume fraction for *k*_p0_*τ*_r_/*g*_eq_ > 1 is given in the Appendix C. In this regime, the deviation has an asymptotic form

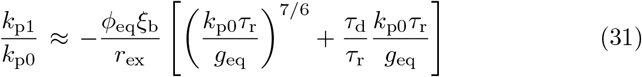

for large polymerization rate *k*_p0_*τ*_r_/*g*_eq_ and ratio *τ*_d_/*τ*_r_, see the cyan solid line in fig. 5**a**. The polymerization rate *k*_p0_ changes not only the polymer volume fraction (compare the solid and broken lines in fig. 3**c**), but also the brush height (see fig. 4) and the diffusion length *l*_d_. To analyze the contribution of the polymer dynamics, we compared the deviation *k*_p1_ derived by using eqs. (7) and (13) and the deviation derived by using the polymer volume fraction of the equilibrium polymer brush. This calculation reveals that the condensation of polymers due to their slow relaxation dynamics decreases the polymerization rate, but only slightly, compare the solid and broken lines in fig. 5**a**. Indeed, the polymer dynamics contributes to the first term of eq. (31) only by the numerical factor of order unity, see Appendix C. The diffusion length *l*_d_ (= *ξ*_c0_) decreases and the brush height *h*_0_ increases with increasing the polymerization rate *k*_p0_. The absolute value of the deviation *k*_p1_ increases with the polymerization rate *k*_p0_ due to the fact that reactant monomers have to travel deeper into the region of large polymer volume fraction in the small length scale (the first term of eq. (31)) and have to travel longer distance (the second term of eq. (31)) with increasing the polymerization rate *k*_p0_.
2. The case of *τ*_d_ < *τ*_r_: The end-grafted polymers are still mushrooms at the crossover between the quasi-equilibrium and non-equilibrium regimes, *k*_p0_*τ*_r_/*g*_eq_ = (*τ*_r_/*τ*_d_)^6/11^, see the magenta line in fig. 4. For *k*_p0_*τ*_r_/*g*_eq_ < (*τ*_r_/*τ*_d_)^6/11^, the distribution of reactant monomers in the entire region of the mushrooms is at the thermodynamic equilibrium, *l*_d_ > *h*_0_, and thus the polymer volume fraction in the mushrooms does not affect the polymerization rate, *k*_p1_ = 0, see the magenta solid line in fig. 5**a**. For (*τ*_r_/*τ*_d_)^6/11^ < *k*_p0_*τ*_r_/*g*_eq_ < (*τ*_r_/*τ*_d_)^15/11^, the polymer volume fraction *ϕ*_p_ in the mushrooms due to the polymer dynamics and the distribution of reactant monomers becomes out of the equilibrium. The deviation *k*_p1_ of polymerization rate due to the polymer volume fraction is negative and its absolute value increases gradually with increasing *k*_p_*τ*_r_/*g*_eq_ through the crossover between the mushroom and the brush regimes at *k*_p0_*τ*_r_/*g*_eq_ = (*τ*_r_/*τ*_d_)^15/11^, see the solid magenta line in fig. 5**a**. The deviation *k*_p1_ of polymerization rate has an asymptotic form

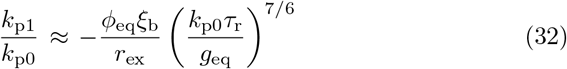

for large polymerization rate *k*_p0_*τ*_r_/*g*_eq_ and small time scale ratio *τ*_d_/*τ*_r_. The condensation of polymers due to their slow relaxation dynamics slightly decreases the polymerization rate *k*_p_, compare the solid and broken magenta lines in fig. 5**a**. The contribution of the polymer dynamics to eq. (32) is only the numerical factor of order unity, see Appendix C. For *τ*_d_ < *τ*_r_, the fact that the diffusion length *l*_d_ decreases with increasing the polymerization rate *k*_p0_ indeed makes the dominant contribution to the suppression of the diffusion of reactant monomers, see also Appendix C. Indeed, if the boundary condition *ϕ*_m_(*r*_c_) = 0 is used instead of eq. (25), the condensation of polymers due to their slow dynamics contributes to the deviation *k*_p1_ more significantly than the case in which eq. (25) is used, compare the magenta solid and broken lines in fig. 5**b**. This implies that the suppression of polymerization rate, eq. (32), is mainly due to the fact that the diffusion length ld decreases with increasing the polymerization rate *k*_p0_. Because of the small height of the brush, the polymer volume fraction in the larger length scales, *z* > *λ*_b_, does not contribute to the polymerization rate significantly even in the brush regime.

**Fig. 5.**
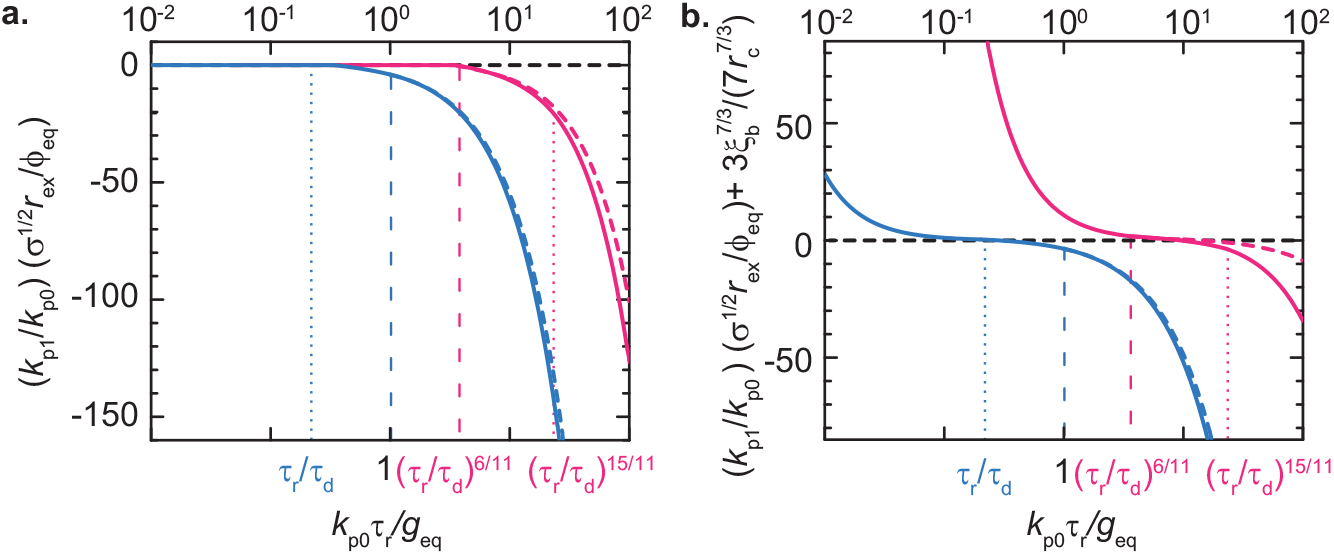
Perturbation analysis: The deviation *k*_p1_ of the polymerization rate due to the polymer volume fraction *ϕ*_p_ in the brush is shown as a function of the polymerization rate *k*_p0_ without the perturbation of *ϕ*_p_ for *τ*_d_ > *τ*_r_ (cyan) and *τ*_d_ < *τ*_r_ (magenta). **a** and **b** are the results of the calculations by using eq. (25) and *ϕ*_m_(*r*_c_) = 0, respectively. The length scale *ξ*_c_ depends on the polymerization rate *k*_p_ and the reaction radius *r*_c_ is a constant. The polymer volume fraction with the polymer dynamics (the solid line in fig. 3**c**) is used to derive the cyan and magenta solid lines. The cyan and magenta broken lines are derived by using the polymer volume fraction at the thermodynamic equilibrium (the broken line in fig. 3**c**). The vertical axis of **b** is shifted from **a** by a constant term 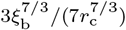 (which is indeed the sole term tha depends on *r*_c_), but **a** and **b** are shown in the same scale. The horizontal axis is logarithmic and the vertical axis is linear. We used *τ*_d_/*τ*_r_ = 5.0 (cyan) and 0.1 (magenta) for the calculation.

## 4 Discussion

We have used the scaling theory to predict the polymer volume fraction and height in a polymer brush inspired by ribosomal RNA transcription. This polymer brush is grown by adding monomers to the grafted-ends of polymers. Our theory predicts that the polymerization kinetics increases the polymer volume fraction in the length scales *ξ*_c_ < *z* < *λ*, see eqs. (3) and (12) for the definition of *ξ*_c_ and *λ*. The polymerization rate decreases as the polymer volume fraction in the brush increases because it suppresses the diffusion of reactant monomers. The deviation *k*_p1_ of polymerization rate due to the polymer volume fraction in the brush is a negative value and its absolute value increases with increasing the polymerization rate *k*_p0_ without the perturbation due to the polymer volume fraction. It is mainly because of the fact that diffusion length *l*_d_ decreases or the brush height increases with increasing the polymerization rate *k*_p0_ and the condensation of polymers due to the slow polymer dynamics plays only a minor role, see eqs. (31) and (32).

For simplicity, we have assumed that the polymerization rate is the diffusion-limited reaction. For the general case, the polymerization rate is calculated by using the rate equation *k*_p_ = *k*_0_*ϕ*_m_(*ξ*_c_)e^*ϕ*_p_(*ξ*,_c_)-*ϕ*_p_(*r*_c_)^ for the boundary condition at *r* = *ξ*_c_, instead of using eq. (25), where *k*_0_ is the rate constant. For the case of the rate-limited reaction, the polymerization rate is trivially *k*_p_ = *k*_0_*ϕ*_m∞_e^-*ϕ*_p_(*r*_c_)^ and the polymer dynamics is not involved as long as *r*_c_ < *ξ*_c_.

Because the polymer brush is inspired by ribosomal RNA transcription, it may be of interest to estimate how much extent the polymerization dynamics affects the structure of nascent pre-rRNAs. The relaxation time of monomers is estimated as *τ*_1_ = *b*^2^/*D*_1_ ≈ 0.3 *μ*s and the elongation rate (per monomer) is 8 s^-1^, see table 1. The number *g*_c_ of monomers in the smallest relaxed blob is thus 5 × 10^4^, corresponding to 600 kb. The putative region, which is localized at the DFC, is 200 b and is much smaller than *g*_c_. Even the total length of pre-rRNA, ≈ 15 kb, is smaller than *g*_c_. This implies that the transcription of pre-rRNAs by Pol I is too slow for the polymer dynamics to be relevant and thus is safe to neglect this effect. This justifies the approach used in our previous theory that neglects the polymer dynamics to predict the multi-phase structure of nucleolus [6].

**Table 1.** Estimate of parameters involved in ribosomal RNA transcription: The Kuhn length of a pre-rRNA is estimated by using the value of single-stranded DNA. The monomer diffusion constant is estimated by 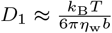, where *κ*_B_*T* (≈ 4 × 10^-21^ J) is the thermal energy and *η*_w_ (≈ 1 mPa · s) is the viscosity of water.

| Parameter | meaning | Value | Ref. |
| --- | --- | --- | --- |
| $b$ | Kuhn length | 4 nm (= 12 b) | [40] |
| $D_1$ | Monomer diffusion constant | $5 \times 10^{-11} \text{ m}^2/\text{s}$ | |
| $v_e$ | Elongation rate | 6 kb/min | [41, 42] |

Our theory predicts that the longest length scale *λ* in which the polymer volume fraction increases due to the polymer dynamics is determined by the relaxation time *τ*_r_ of a subchain in the equilibrium blob, which is much smaller than the relaxation time of the whole chain. One may wonder why the relaxation of a polymer in a brush is faster than the relaxation of a free polymer in a solution. Indeed, it is not the case. In the thermodynamic equilibrium, a polymer brush is viewed as a melt of equilibrium blobs [9]. The equilibrium blobs in a chain show the lateral fluctuation with the magnitude *δR*_eq_ = *ξ*_b_(*N*/*g*_eq_)^1/2^. The relaxation time of the whole chain is the time scale of the lateral excursion of equilibrium blobs. We thus expect that the magnitude of the lateral fluctuation is *ξ*_b_ for the length scales *ξ*_b_ < *z* < *λ* and increases with the distance *z* for *z* > *λ* because the distance *z* reflects the time *t* after the addition of monomers in the blobs, see fig. 6**a**. Because of the connectivity of blobs, the blobs show the Rouse dynamics. The magnitude of the fluctuation for *z* > *λ* is thus *δR* = *ξ*_b_(*t/τ*_r_)^1/4^ (= *ξ*_b_(*z/λ*)^1/4^), see fig. **6a** (or **b** represented as a function of the number *n* of monomers from the grafted ends). This implies that polymers in a dynamic brush show smaller lateral excursion than polymers in an equilibrium brush. The relaxation time of the whole chain does not affect the polymer volume fraction *ϕ*_p_ in the brush.

**Fig. 6.**
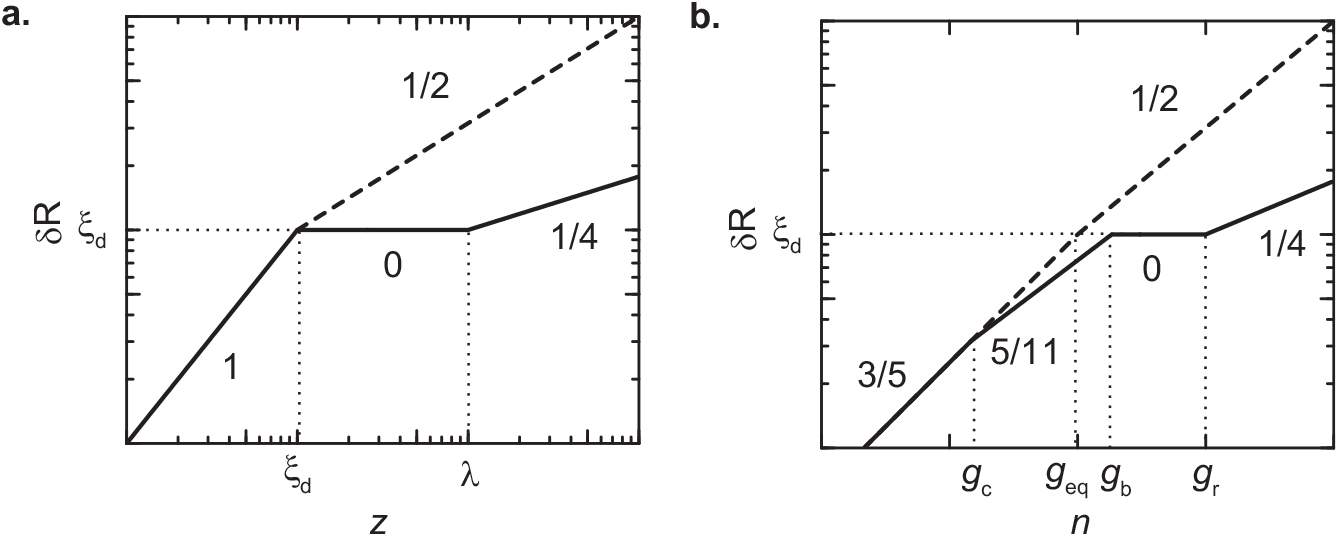
Magnitude of lateral fluctuation: The magnitude *δR* of the lateral fluctuation is shown as a function of the distance *z* from the surface (**a**) and the number *n* of monomers from the grafted ends (**b**) of polymers in a non-equilibrium polymer brush (solid) and an equilibrium brush (broken).

Inspired by the transcription of pre-rRNAs, we have proposed a brush of polymers, where monomers are added to their grafted-ends with a constant rate. This brush is different from a brush of polymers, where monomers are added to their free-ends [33, 34], such as the case of free radical polymerization from initiators grafted at a surface. However, the difference is only significant when the polymerization kinetics is fast or polymers in the brush are very long. As such, one does not have to worry about the slow relaxation dynamics of polymers in the context of the transcription of pre-rRNAs. We anticipate that our theory may be extended to polymerization reactions in other biological contexts, where the slow relaxation dynamics of polymers is relevant, and/or may motivate polymer chemists to design a polymer brush by taking advantage of their slow relaxation dynamics and polymerization kinetics.

## Acknowledgments

This work was supported by JSPS KAKENHI Grant Number 20H05934 and 21K03479. The authors acknowledge valuable comments by Michael Rubinstein (Duke Univ.).

## Appendix A Chemical potential of reactant monomers

The mean field free energy of the subchain (composed of g monomers) in a relaxed blob (of the size of *ξ*) has the form

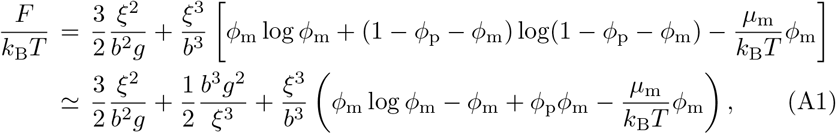

where *ϕ*_m_ and *μ*_m_ are the volume fraction and chemical potential of reactant monomers. *k*_B_ is the Boltzmann constant and *T* is the absolute temperature. In the first form of eq. (A1), we take into account the free energy due to the entropic elasticity of the subchain (the first term) and the mixing entropy of solvent molecules and reactant monomers (the second term). We assumed that the solvent is athermal both for polymers and reactant monomers because they have the same chemistry. The last form of eq. (A1) is derived by expanding the free energy in a power series of *ϕ*_m_ and *ϕ*_p_ and neglected the higher order terms.

The minimization of eq. (A1) with respect to *ϕ*_m_ leads to the form of the chemical potential

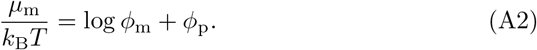

The minimization of the free energy with respect to the size *ξ* leads to the form

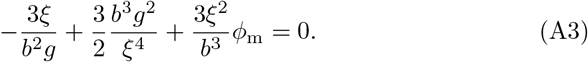

Eq. (A3) represents the balance of the elastic force of the subchain (first term), the force due to the excluded volume interaction between monomers in the subchain (second term), and the force due to the osmotic pressure of reactant monomers (third term). For simplicity, we here treat cases in which the osmotic pressure of reactant monomers is small and thus does not affect the size of relaxed blobs. In such cases, the size of a relaxed blob is *ξ* ≈ *bg*^3/5^ and thus the polymer volume fraction *ϕ*_p_ derived in the previous section can be used as it is.

## Appendix B Volume fraction of reactant monomers

The monomer volume fraction *ϕ*_m_(*z*) for *z* > *ξ*_b_ has the form

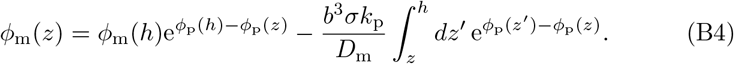

Eq. (B4) is derived by using eqs. (21), (20), and (22) with the boundary condition, eq. (24). The monomer volume fraction *ϕ*_m_(*r*) for *r* < *ξ*_b_ has the form

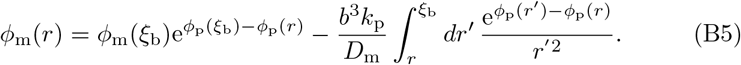

Eq. (B5) is derived by using eqs. (21), (20), and (23). Because the polymer volume fraction *ϕ*_p_ is continuous at *z* = *ξ*_b_, the monomer volume fraction *ϕ*_m_ is also continuous there. In the mushroom regime, the monomer volume fraction *ϕ*_m_(*r*) is derived by replacing *ξ*_b_ to *h*.

## Appendix C Deviation of polymerization rate by polymer volume fraction

We here summarize the deviation *k*_p1_ of polymerization rate by the polymers in the brush as a function of *p* (= *k*_p0_*τ*_r_/*g*_eq_):

i) The case of *τ*_d_ > *τ*_r_.
1) Brush regime - with polymer dynamics:

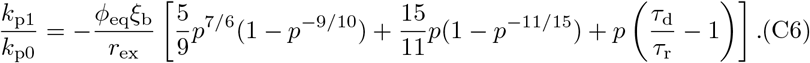
- without polymer dynamics:

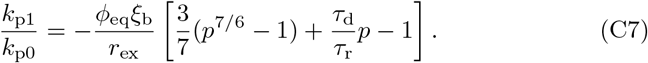
ii) The case of *τ*_d_ < *τ*_r_.
2) The mushroom regime - with polymer dynamics:

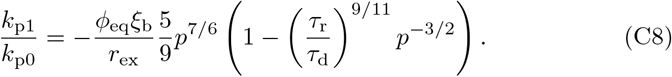
- without polymer dynamics:

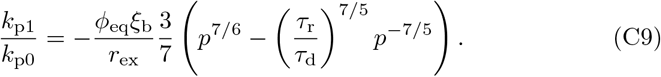
3) Brush regime - with polymer dynamics

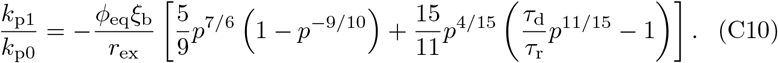
-without polymer dynamics

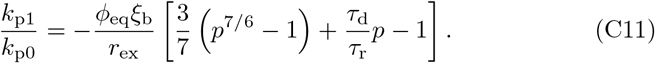

If the boundary condition *ϕ*_m_(*r*_c_) = 0 is used (instead of eq. (25)), the lower boundary *ξ*_c0_ of the integral in the second term of eq. (29) should be replaced to *r*_c_. In this case, the deviation *k*_p1_ is calculated by using

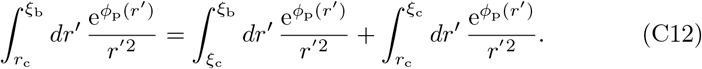

The deviation *k*_p1_ including the first term of eq. (C12) has been already derived in eqs. (C6) - (C11). The deviation *δk*_p1_ due to the second term of eq. (C12) is

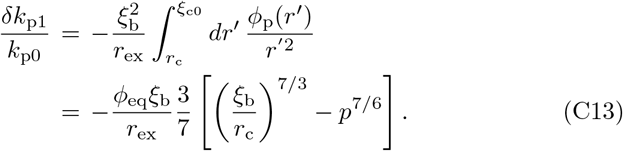

The deviation for the new boundary condition is *k*_p1_ + *δk*_p1_. We note that the polymer dynamics does not affect the polymer volume fraction *ϕ*_p_ in the length scales *r*_c_ < *r* < *ξ*_c0_. We thus used the polymer volume fraction for the equilibrium to derive eq. (C13).

## Notes

### Competing Interest Statement

The authors have declared no competing interest.

